# Replication-dependent organisation constrains positioning of long DNA repeats in bacterial genomes

**DOI:** 10.1101/2022.03.16.484558

**Authors:** Nitish Malhotra, Aswin Sai Narain Seshasayee

## Abstract

Bacterial genome organisation is primarily driven by chromosome replication from a single origin of replication. However chromosomal rearrangements, which can disrupt such organisation, are inevitable in nature. Long DNA repeats are major players mediating rearrangements, large and small, via homologous recombination. Since changes to genome organisation affect bacterial fitness - and more so in fast-growing than slow-growing bacteria - and are under selection, it is reasonable to expect that genomic positioning of long DNA repeats is also under selection. To test this, we identified identical DNA repeats of at least 100 base pairs across ~6,000 bacterial genomes and compared their distribution in fast and slow growing bacteria. We find that long identical DNA repeats are distributed in a non-random manner across bacterial genomes. Their distribution differs in their overall number, orientation and proximity to the origin of replication, in fast and slow growing bacteria. We show that their positioning - which might arise from a combination of the processes that produce repeats and selection on rearrangements that recombination between repeat elements might cause - permits minimum disruption to the replication-dependent genome organisation of bacteria, thus acting as a major constraint.

## Introduction

Most bacterial genomes consist of a single circular chromosome. Similar to the genomes of all living organisms, the bacterial genome is condensed and organised inside the cell. Despite sharing different strategies of gene organisation with eukaryotes (Lawrence 2002), unlike eukaryotes, bacterial genome organisation is primarily driven by chromosome replication (Rocha 2004).

Chromosome replication in bacteria begins at a single locus called the origin of replication; oriC; and terminates diametrically opposite at the terminus of replication, ter (Duggin & Bell 2009). The movement of the replisome from oriC to ter is bidirectional, creating replichores on both sides of the oriC. The average doubling time of fast-growing bacteria such as *Escherichia coli* is less than the time required to replicate its chromosome (Cooper & Helmstetter 1968). To compensate for the lag between chromosome replication time and division time, a new replication cycle begins at oriC even before the previous one ends at ter. Consequently, at any point of time during replication, the copy number of regions near oriC is higher than those near ter resulting in an oriC-ter dosage gradient. This dosage gradient can be as high as 8:1 in *E. coli*. Even in slower-growing bacteria in which DNA replication occupies a substantial portion of the cell cycle, a 2:1 dosage gradient between oriC and ter will be common. As a possible consequence of selection arising from this oriC-ter gene dosage gradient, highly expressed genes - primarily those coding for the translation machinery - are found proximal to the oriC locus whereas stress response genes and horizontally acquired genes are localised proximal to ter (Rocha 2004; Couturier & Rocha 2006)

In addition to the dosage gradient observed, the genomic content in bacteria is differentially distributed across strands and replichores formed during replication (Rocha 2004). Nucleotide composition in the leading strand is skewed towards G and T whereas the region around ter is A-T rich. The leading strand is also abundant in DNA motifs involved in recombination-mediated repair viz. Chi and chromosomal segregation viz. KOPS. Furthermore, it encodes more genes than the lagging strand, and is in particular enriched for essential and highly expressed genes. This is often attributed to the detrimental effect of head-on collisions between the DNA polymerase and the RNA polymerase, which are more likely to happen during transcription of highly expressed genes encoded on the lagging strand (Rocha 2003, 2004, 2008).

Despite organisational features of the genome being shared across bacteria (Couturier & Rocha 2006; Khedkar & Seshasayee 2016; Tamames 2001), the events that generate chromosome variations are inevitable in nature and large rearrangements are one of the major contributors to these variations. Chromosome rearrangements like deletions, duplications and inversions are caused by cellular processes like homologous recombination and transpositions under different genetic and environmental conditions (Bishop 2000; Veetil et al. 2020; Srinivasan et al. 2015; Sonti & Roth 1989). DNA repeats are one of the major players mediating such events (Achaz et al. 2003; Bi & Liu 1996; Lambert et al. 1999; Treangen et al. 2009). DNA repeats or duplicated stretches of DNA range from dinucleotide to thousands of nucleotides and are abundant in bacterial genomes (Treangen et al. 2009). DNA recombination between repeats results in structural variations and is linearly dependent on substrate length (Shen & Huang 1985). ‘Long’ DNA repeats, i.e., DNA repeats of length of at least 100 nucleotides can lead to intra-chromosomal rearrangements by acting as substrates to the bacterial recombination machinery (Treangen et al. 2009; Shen & Huang 1985).

The type of rearrangements mediated by long DNA repeats depend on the relative orientation of the repeat pairs. Direct repeats - i.e., repeat pairs present in the same orientation - result in duplication or deletion of the genomic region flanked by them. On the other hand, inverted repeat pairs - i.e., repeat pairs that are positioned in the genome in opposite orientation - lead to inversion of the genomic region. Deletions lead to the removal of the chromosomal region thus resulting in gene loss whereas duplications lead to doubling of the genomic segment consequently increasing the copy number and presumably the expression levels of the affected genes (Skovgaard et al. 2011; Srinivasan et al. 2015; Sonti & Roth 1989; Straus & Hoffmann 1975). On the other hand, inversions caused by inverted repeat pairs flip the repeat-flanked region thus reversing its orientation and can cause detrimental head-on collisions between the two polymerases. Large inversions also result in a significant disruption of gene dosage gradient, affecting fitness particularly in conditions supporting fast growth (Srivatsan et al. 2010). Taken together, repeat mediated rearrangements disrupt the bacterial genome organisation by altering the dosage and orientation of genes.

Previous studies have reported such events of bacterial rearrangements under different stresses. The rearrangements observed were associated with repeat elements like insertion sequences, and were in turn advantageous or disadvantageous in different environments (Veetil et al. 2020; Repar et al. 2017; Srinivasan et al. 2015; Sonti & Roth 1989; Adler et al. 2014; Maharjan et al. 2013). Since these repeats-associated changes in genome organisation play a role in affecting fitness, there might be selection on the positioning of such repeats on the chromosome. Studies indicating non-random genomic distribution of repeats or landscapes of chromosomal rearrangements suggest chromosomal composition, relative position with respect to origin of replication, or pathogenicity as constraints on genomic presence of such repeats (Rocha et al. 1999; Repar & Warnecke 2017).

In this study we used comparative genomics to investigate the association between replication-dependent genome organisation and long DNA repeats. Through this work, we asked the following questions on the genomic distribution of long DNA repeats, 1) Are long repeats present randomly across genomes? 2) Does their distribution reflect selection imposed by the nature of structural variation they mediate? 3) How does their genomic arrangement vary across bacteria with different growth rates? Using ~6000 bacterial genomes across different genera and classes of bacteria representing fast and slow growing bacteria, we found that long identical DNA repeats are distributed non-randomly across bacterial genomes. The genomic distribution of these repeats differs in number, orientation and oriC-proximity in fast and slow growing bacteria. We found that despite differential genomic distribution of such repeats the repeat pairs are present in such a way that limits disruption to the replisome. This arrangement of repeat pairs is more prominent in fast growing bacteria where the effect of replication machinery is seen to a greater extent. Taken together, our study identifies bacterial growth as one of the constraints to genomic organisation of long DNA repeats in bacterial genomes.

## Material and Methods

### Data

DNA sequence files (.fna) and genomic feature files (.gff) of 6387 completely sequenced (as of april 2019) bacterial genomes were downloaded from RefSeq database (O’Leary et al. 2016) at NCBI ftp website. Only the main chromosome of a genome was used for this study. The classification of these genomes into phyla, class, genera and species was done on the basis of their information available on sequence headers and Kegg classification of NCBI genomes (https://www.genome.jp/kegg/docs/cmp_prok.html). These genomes had at least one of their species representatives with a single predicted origin of replication in the DoriC database(Gao & Zhang 2007) (DoriC 6.5, http://tubic.tju.edu.cn/doric/public/index.php).

### Classification of bacteria based on growth

Bacterial genomes were classified into two categories: fast growing and slow growing by calculating a parameter called Rf factor (Couturier & Rocha 2006; Khedkar & Seshasayee 2016). It was defined as the ratio of estimated time taken for a full round of chromosome replication (Tr) and minimum doubling time (Td). Tr was calculated by taking the ratio of half the genome size to 600 nt/sec which is an average speed of DNA replication(Reyes-Lamothe et al. 2008; Milo et al. 2009). For Tr, genome sizes were defined as string length of the sequence in .fna files using in house python script, while Td was estimated by rDNA copy number and doubling time of the bacteria as mentioned in (Freilich et al. 2009; Couturier & Rocha 2006; Khedkar & Seshasayee 2016)

Bacteria with Rf >1 are expected to initiate replication more than once per cell cycle on average which is true for fast growing bacteria. On the other hand, slow growing bacteria have Rf <= 1.

### Identification of long intra-chromosomal DNA repeats and associated genomic regions

Identical long repeats of length at least 100 base pairs were identified by the repeat-match algorithm of the MUMmer software using -n 100 option(Delcher 2002; Kurtz et al. 2004). The positions and orientation of the repeat pairs were extracted using an in-house python script. Repeat pairs were classified as direct pairs when both the repeat partners are in the same orientation, i.e., the positions of the exact matches in the mummer output are on the same strand and inverted pairs when it was otherwise.

Genomic coordinates of CDS and rDNA regions were extracted from the .gff files. IS elements, prophages and horizontally acquired regions were identified using ISEScan (Xie & Tang 2017), Phaster (Arndt et al. 2016), and AlienHunter (Vernikos & Parkhill 2006)) respectively. To identify the genomic regions associated with repeats, overlap between the identified repeats and these regions were calculated by comparing the respective genomic coordinates. A repeat was reported to be associated with these regions if there was an overlap of even a single nucleotide base with the respective region. Repeat pairs in which any of the repeat partners had an overlap with rDNA regions were removed from further analysis. This led us to have 6340 genomes with at least one repeat pair.

### Association of inversions with inverted repeat elements

For every species in our dataset, a reference genome was chosen randomly and other genomes were used as a query for predicting inter-genomic inversions. To predict large chromosomal inversions of at least 1kb size, nucmer -l 1000 function of MUMmer (Delcher 2002; Kurtz et al. 2004) package was used. Using the orientation information in the output file, large inter-genomic inversions were found. Their positions were parsed using in-house python scripts and compared with the positions of intra-chromosomal long inverted repeats. Only those genomes were chosen for the analysis which had at least 30 distinct genomic positions with repeats in the genome and at least one predicted inversion with respect to the reference. This left us with 3730 genomes for this analysis.

For every inversion detected in the genome with respect to the reference, distance (in bases) was calculated between the nearest repeat (as part of inverted repeat pairs identified) and the inversion observed. This gave us 4 different distance values; from 5’ and 3’ sides in the query and its reference; and amongst which the shortest distance was used to compare the distances for this particular analysis.

### Calculation of relative position of repeats with respect to oriC

For every bacterial species, the oriC of a representative genome in the DoriC database was used as a reference and blastn (Altschul et al.) was performed to predict oriC sequences in the rest of the genomes of that species. Blast hits with the highest score and with at least 90% identity was chosen as the oriC sequence of that genome. For every genome with predicted oriC, the shortest distance between the repeat and the oriC was calculated thus providing the relative position of repeats with respect to origin of replication. The relative position was then normalised to 1000 bp i.e., divided by genome size and then multiplied by 1000 to make it comparable across genomes.

### Calculation of genomic distribution of repeats

To understand the genomic distribution of repeats, every genome was divided into equal parts of 2, 4,and 20 bins centred around the oriC-ter axis. Ori bin was centred at oriC while ter bin was referred to as the region diametrically opposite to the Ori bin. Left and right bin(s) were classified as regions which were on the left replichore or upstream of oriC and right replichore or downstream of oriC respectively. This is similar to the approach taken in Khedkar & Seshasayee 2016.

In order to calculate bin level enrichment, repeats were counted for every bin using their normalised positions and were divided by total number of repeats in the genome. These genomes (5970 genomes)had at least 30 distinct genomic positions with repeats. While comparing the enrichment between fast-growing and slow-growing bacteria, the distributions of medians of proportions across bins were made across two groups and compared using F-test. Similar analysis was performed for calculating intra-quadrant repeat enrichment.

### Calculation of distances and symmetry between repeat pairs

For intra-replichore repeat pairs, distance between repeats was calculated as the absolute difference in normalised position of repeat pairs. And symmetry in inter-replichore repeat pairs was calculated as 1 - absolute difference in the normalised relative position of the repeat units with respect to oriC, similar to Khedkar & Seshasayee 2016. These values ranged from 0 to 1 with 0 being most symmetric around the ori-ter axis while 1 being least symmetric, i.e., repeat pair is on the axis itself. For this analysis, the genomes that had at least 30 distinct positions of repeats involved in direct and inverted repeat pairs were taken. A total of 4165 genomes were used in this analysis.

### Null models and comparisons across parameters

To test the statistical significance of the observed values of the parameters in all our analysis, we compared the observed dataset relevant to that analysis with that of the null model generated through randomisation. The underlying assumptions of all comparisons were that individual repeat elements are independent of each other and there is no other constraint than bacterial growth. To consider phylogenetic dependency of the genomes, we show major observations at phyla level and also after removing redundancy in species. All the comparisons and their corresponding null models taken are explained in the relevant section.

## Results

### Repeat density is not correlated with bacterial genome size or growth

The premise of this study is that long DNA repeats i.e., DNA repeats of at least 100 bp length are capable of intra-chromosomal recombination leading to a variety of structural variations (Shen & Huang 1985). These variations have the potential to alter genome organisation and can be beneficial or deleterious in different conditions. Consequently, this might impose selection on where such repeats are positioned on the chromosome.

We identified intra-chromosomal identical repeats whose repeating units are at least 100 base pairs (bp) in length using the MUMmer (Delcher 2002; Kurtz et al. 2004) package across bacterial genomes obtained from the NCBI RefSeq database (O’Leary et al. 2016). These genomes represent 653 bacterial species from 462 genera and 27 phyla.

~78% of DNA repeats overlapped coding sequences (CDS) and ~11% had an overlap with rDNA regions. Additionally, an average of ~60% of the repeats were part of horizontally acquired regions. Insertion Sequences (IS) elements and prophages constitute ~23 % and ~2% of the repeats respectively (Figure 1a). Note that these regions can be overlapping with each other and therefore the percentages may not sum up to 100. Since in this study we investigate the genomic distribution of DNA repeats, we specifically removed repeat pairs that had an overlap with rDNA regions (i.e., an average ~11% of repeats). This is because highly expressed rDNA loci are often present closer to oriC and as a consequence can contribute to unwanted bias in our study. This left us with a median of 120 distinct repeats ranging from a minimum of 1 repeat to 9217 repeats per genomes.

**Figure 1.**
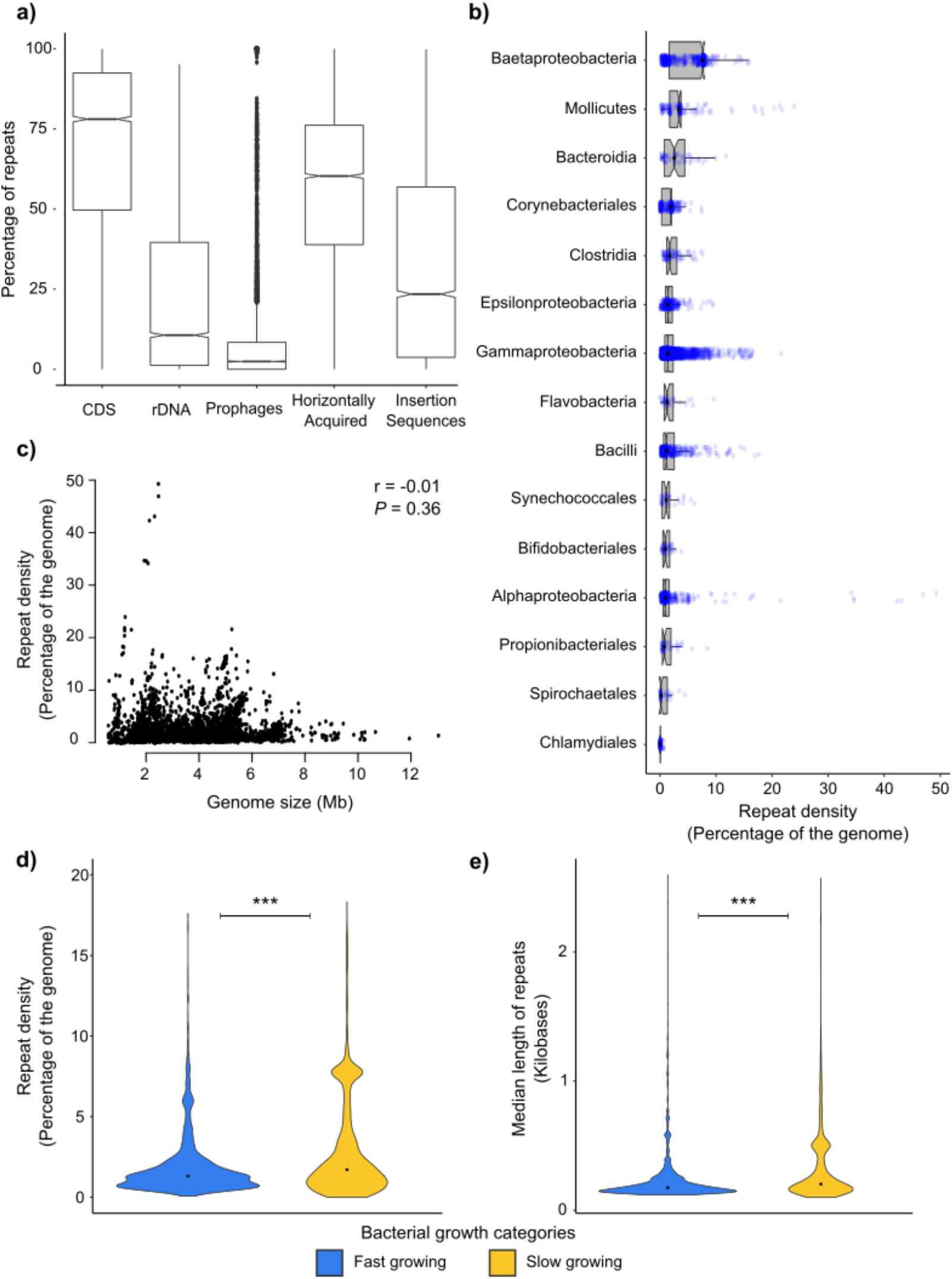
(a) Boxplots of percentage of repeats overlapping with annotated CDS, rDNA regions and predicted prophages (using Phaster), horizontally acquired regions (using Alien Hunter) and insertions sequences (ISEScan). (b) Boxplots of repeat densities across classes of bacteria with blue jitter showing the distribution. Only classes with at least 30 genomes are shown here. (c) Scatter plot showing relation between repeat density and genome size (Megabases; Mb). The correlation is calculated by Pearson’s correlation using cor.test() function in R. (Total number of genomes = 6340). (d) Violin plots showing repeat density distribution in fast growing and slow growing bacteria. Wilcoxon’s rank-sum test is used to calculate the significance in the differences of two medians. (e) Violin plots showing distribution of median length of repeats in fast growing and slow growing bacteria. Wilcoxon’s rank-sum test is used to calculate the significance in the differences of two medians. (n.s.:*P*> = 10^−2^, * : *P* < 10^−2^, ** : *P*< 10^−4^, *** : *P*< 10^−6^)

The repeat density, defined by the proportion of the repetitive genome, ranged from ~0.005% in *Advenella* to ~49% in *Orientia* with a median of ~1.5%. This density varied across different classes of bacteria (Figure 1b) and did not correlate (Pearson’s correlation coefficient; r = −0.011, *P*= 0.36, N = 6340) with genome size (Figure 1c). We observed the same even after removing the redundancy at the species as well as genera level (Pearson’s correlation coefficient; r = −0.043, P= 0.26, N= 653) (Figure S1). This is contrary to an earlier study done on a smaller set (N = 53) of genomes by (Rocha et al. 1999)which found a negative correlation between repeat density and bacterial genome size.

Bacterial cells spend a substantial portion of their cell cycles replicating the chromosome. In fast growing organisms in which chromosome replication takes longer than the average population doubling time, there are multiple replication cycles ongoing at anytime. This establishes an oriC-ter dosage gradient. Even in slow-growing bacteria, there will be a gradient for the proportion of the cell cycle during which replication is going on. However, the oriC-ter gradient will be steeper in fast-growing bacteria than in slower-growing ones. In order to incorporate this effect of growth rate on the replicative structure in our analysis we classified bacteria into fast- or slow-growing organisms and compared the genomic distribution of repeats between them. The bacterial growth categorisation is based on a rough estimate of the oriC-ter gradient determined by (Couturier & Rocha 2006), defined as the number of replication initiations per cell cycle. Fast-growing bacteria are defined as those with Rf > 1; where the chromosome replication time is greater than the average doubling time, indicating a steep replication-dependent gene dosage gradient. In contrast, slow-growing bacteria have Rf <= 1. In this study we have identified 2337 fast-growing and 4003 slow-growing bacteria with at least one repeat pair in their genomes. In this categorisation, fast-growing bacteria span 6 phyla and 16 classes while slow-growing bacteria span 27 phyla and 75 classes of bacteria (Supplementary File available at https://doi.org/10.6084/m9.figshare.19367048).

We note that the repeat density of fast-growing bacterial genomes is significantly lower than that of slow growing bacteria (Wilcoxon rank sum test; P < 10^−10^) (Figure 1d). We further observed that the median length of repeats in fast growing bacteria is also lower than slow growing bacteria (Wilcoxon rank sum test; P < 10^−10^) (Figure 1e). These suggest that repeat-mediated recombination events may be selected against particularly in fast-growing bacteria.

### Inverted repeats are underrepresented in bacterial genomes

Following the identification of long DNA repeats in bacterial genomes, we studied the distribution of repeats responsible for specific kinds of structural variation. The nature of structural variation caused by repeat pairs depends on their relative orientation. Repeat pairs can be classified as direct if both repeats in a pair are oriented in the same direction or inverted if they are oriented opposite to each other. Direct repeats can lead to duplications and deletions whereas inverted repeats can cause inversions of the region flanked by them.

In our dataset, we found that ~79% (4984/6340) genomes have a lower proportion of inverted repeats than direct repeats (i.e., number of inverted repeat pairs/ number of direct repeat pairs <1). This is consistent with the earlier report by (Achaz et al. 2003). Similar to their report, we observed that this lower proportion of inverted repeats is more prominent in genomes with lower number of repeat pairs. This proportion approaches 1 with increase in repeat pairs (Figure 2a). To understand that if this observed proportion is just by random chance, we generated a null distribution by allocating an orientation to each repeat element in a genome randomly. While creating the null distribution, we keep the number of pairs the same as that in the observed set. We further calculated the count of direct and inverted repeat pairs for that genome for 1000 such random allocations. For every genome, we applied a z-statistic on the observed count of direct and inverted repeats and calculated the P-value associated with the count. We found that ~46 % (2903/6340) of the genomes had a significantly lower number of inverted repeat pairs (z<0, P <= 0.01) while just ~2% (106/6340) of the genomes had significantly higher proportion inverted repeat pairs (z >0, P <= 0.01). Slow growing bacteria (median inverted repeat proportion; 0.62) encode a relatively lower proportion of inverted repeat pairs than fast growing organisms (median inverted repeat proportion; 0.74) (Wilcoxon rank sum test; P < 10^−10^) (Figure 2b).

**Figure 2.**
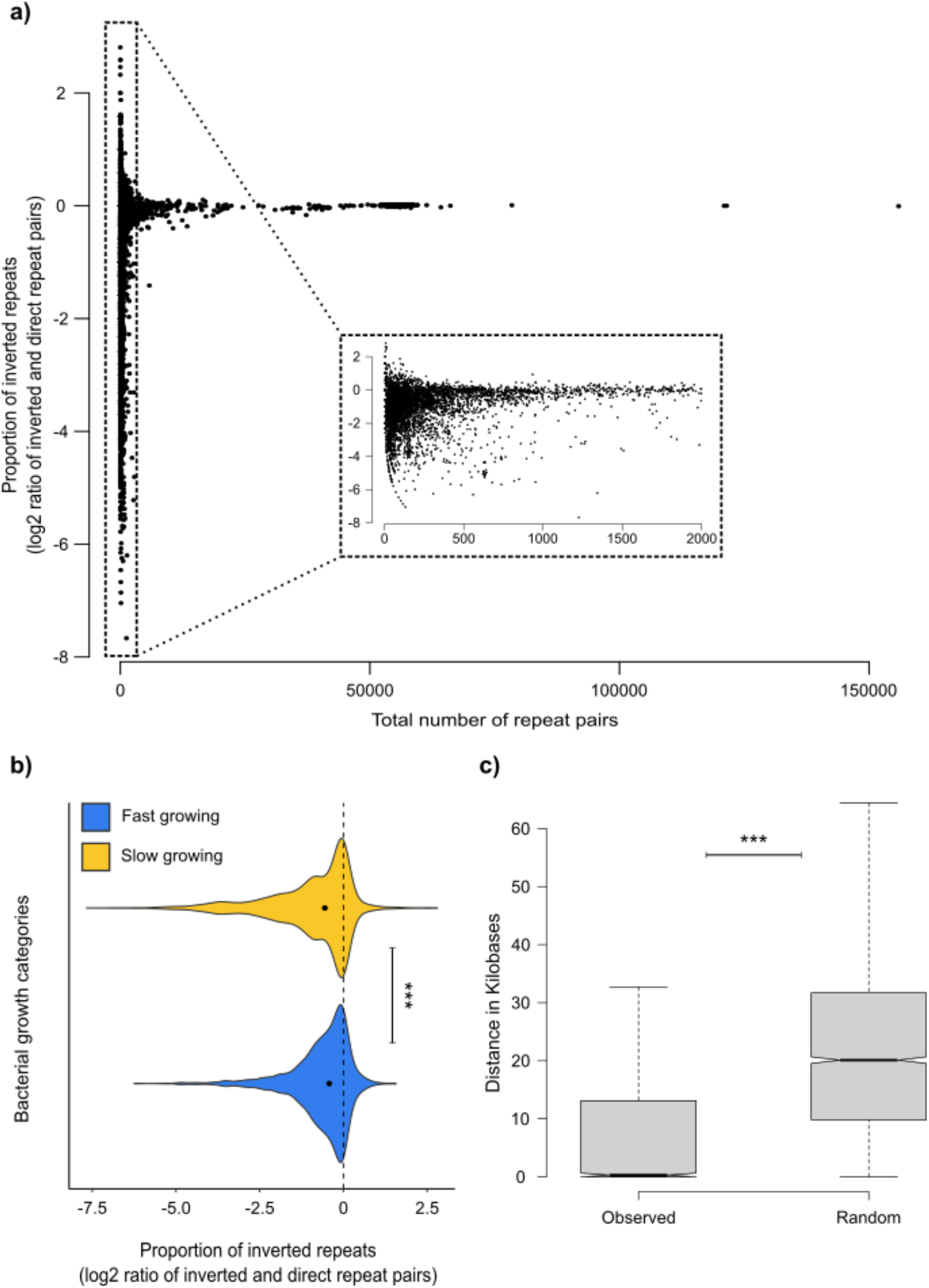
(a) Scatter plot showing relation between proportion of inverted repeat pairs shown as log2 (number of inverted repeat pairs / number of direct repeat pairs). The inset shows a zoomed in version from 0-2000 repeat pairs where inverted repeat pairs are underrepresented and have not reached closer to that of direct repeat pairs. (b) Violin plots showing distribution of log2 proportion of inverted repeat pairs in fast growing and slow growing bacteria. Wilcoxon’s ranksum test is used to calculate the significance in the differences of two medians. (c) Boxplots of the median of minimum distances (in Kilobases; Kb) of the inverted repeats with the genomic inversion in the query sequence with respect to the reference. For every genome, there is one point in each of observed and random set. Wilcoxon’s signed-rank test is used to calculate the significance in the differences of two medians. (n.s. :*P*>= 10^−2^, * : *P* < 10^−2^, ** : *P*< 10^−4^, *** : *P*< 10^−6^)

The presence of significantly lower inverted repeats suggests that the process by which repeats are generated is more likely to produce direct repeats (see Discussion) and / or that selection against inversions in bacterial genomes across genera, the two explanations not being mutually exclusive.

### Repeats are associated with large genomic rearrangements in bacteria

To test if the role of repeats in causing chromosomal rearrangements can be seen in extant genomes, we analysed inverted repeats and their association with large (> 1kb) chromosomal inversions. We specifically selected inversions for this analysis because large inversions are relatively more stable than large deletions and duplications (Straus & Hoffmann 1975; Adler et al. 2014), and are relatively simpler to detect in the assembled genomes. For each species in our dataset, we randomly chose a reference genome and compared all genomes in that species with the reference using MUMmer (Delcher 2002; Kurtz et al. 2004). We extracted genomic positions of large inversions (>= 1Kb) with respect to the reference and then calculated the distance of the nearest repeat element flanking that inverted region in both the reference and the compared genome. We further selected the shortest distance amongst the distances obtained in either the reference or the compared genome and then tested the null hypothesis that the shortest distances between inversions and the nearest inverted repeat element are not significantly different from that obtained if repeats were randomly distributed. The apriori condition for a repeat to be considered was that it should form an inverted pair in the same genome i.e., another instance of the repeat should be present in the same genome but with opposite orientation. The underlying assumption for this analysis is that repeats responsible for the rearrangement in that genome are found in close proximity to the site of inversion. We generated a null model for every bacterial genome by assigning inverted repeat elements to random positions on the genome. To test our hypothesis, we compared distributions of median shortest distances observed in the genomes to the distribution of median distances obtained from 1000 iterations using a Wilcoxon signed rank test (paired). Using the null model thus generated and the statistical tests performed, we rejected the null hypothesis (P<10^−10^) and found that the average distances in the observed dataset are significantly lower than expected had the null hypothesis been true (Figure 2c).

### Long DNA repeats are distributed non randomly across bacterial genomes

Having established that repeats are indeed associated with structural variations, we set out to understand whether there are any constraints on the genomic positions of such repeats. To test the null hypothesis that repeats are positioned randomly on the chromosome, we considered genomes with at least 30 distinct genomic positions with repeats (5970 genomes). For every genome, we calculated the distances between adjacent repeat elements and compared its distribution with the distribution obtained on randomising their genomic positions. The comparisons were done using Kolmogorov Smirnov two-sample test (KS test) for 1000 random iterations followed by Bonferroni test for multiple correction resulting in 1000 P values for every genome. A genome is called to have non-random distribution of repeats if the distance distribution of repeats shows significant deviation from the random distribution i.e., at least 90% of the iterations of the comparison mentioned has a P <= 0.01. We observed that in ~83% (4977/5970) of the bacteria, the genomic distribution of long DNA repeats is non-random. To reduce the effect of strict identity threshold (100%) to predict repeats and overrepresentation of repeats in our analysis due to possible overlaps between proximal repeats, we merged repeats that were separated by <= 100 bases to one repeat. This led us with 5520 genomes with at least 30 repeats. Following merging, we found that only ~38 % (2086/5520) of the bacteria show non-random genomic distribution of repeats. This deviation of genomic distribution of repeats from random can be seen across classes of bacteria with varying degrees. (Figure 3a and 3b). Note here that the stringent P-value thresholds used here might somewhat compromise sensitivity in rejecting the null hypothesis.

**Figure 3.**
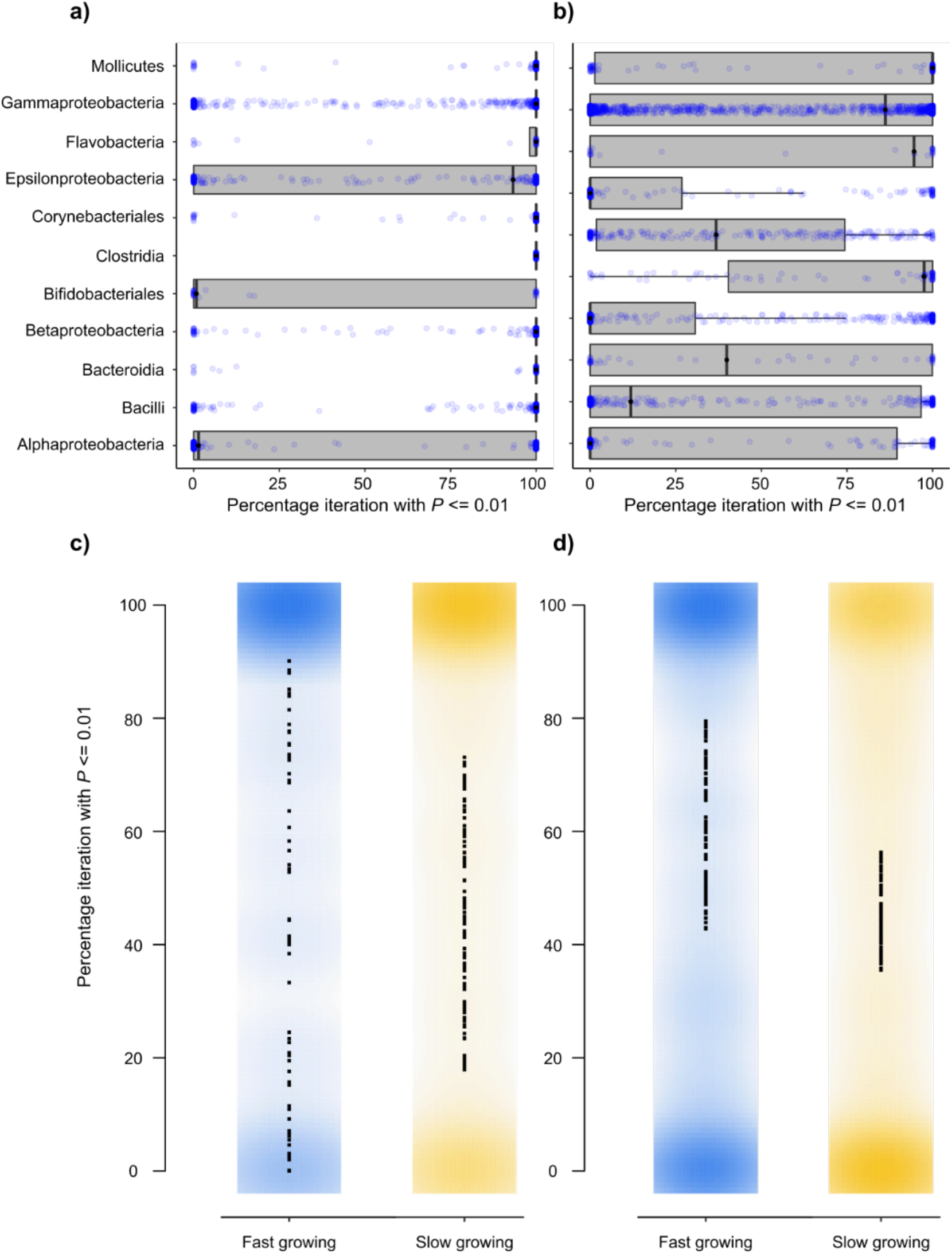
Boxplots of percentage iterations with *P* <= 0.01 obtained to check randomness in repeat distribution of repeats (a) without merging the overlapping repeats and (b) after merging overlapping repeats with blue jitter showing the distribution. Only classes with at least 30 genomes are shown here. Smooth scatter plot of percentage iterations with *P* <= 0.01 obtained to check randomness in repeat distribution in fast and slow growing bacteria (c) without merging the overlapping repeats and (d) after merging overlapping repeats. The *P* values in all the plots were calculated using KS test between the observed distribution of distances between repeats to that of random for 1000 iterations. The *P* values was adjusted using Bonferroni correction for multiple correction.

The presence of long repeats in a non-random manner at least in a subset of genomes suggests that there is a constraint on the genome-wide organisation of these repeats in bacterial genomes. Since bacterial genome organisation is majorly driven by replication, we further investigated the constraints imposed by replication on the distribution of repeats. On comparing the number of genomes showing non-random repeat distribution, we found that ~90% (2113/2336) of fast-growing bacteria and ~78% (2864/3634) of slow-growing bacteria have non-random genomic distribution of repeats (>= 90% iterations with P<= 0.01). On merging the proximal repeats, the number changes to ~50% (1169/2326) fast-growing bacteria and ~29% (917/3194) slow growing bacteria. (Figure 3c and 3d).

To further understand the difference in non-randomness in genomic distribution of repeats in fast-growing and slow-growing bacteria, we investigated if there are regions in the chromosome that are enriched with repeats. We divided each genome into 20 equally-sized bins taking oriC and ter as reference positions and then calculated the proportion of repeats in each bin as the percentage of bin covered with repeats (Figure 4a). We observed that the repeat distribution across bins varied in fast and slow growing bacteria (Figure 4b). On comparing the distributions of median proportion of repeats across bins, we found that the variance of the repeat proportion across bins was significantly low (F-test; P= 0.0005) in slow growing bacterial genomes as compared to fast-growing bacterial genomes (Figure 4c). This strengthens the implication of replication derived constraints on genomic distribution of repeats and suggests a greater constraint in the position of repeats in fast-growing organisms.

**Figure 4.**
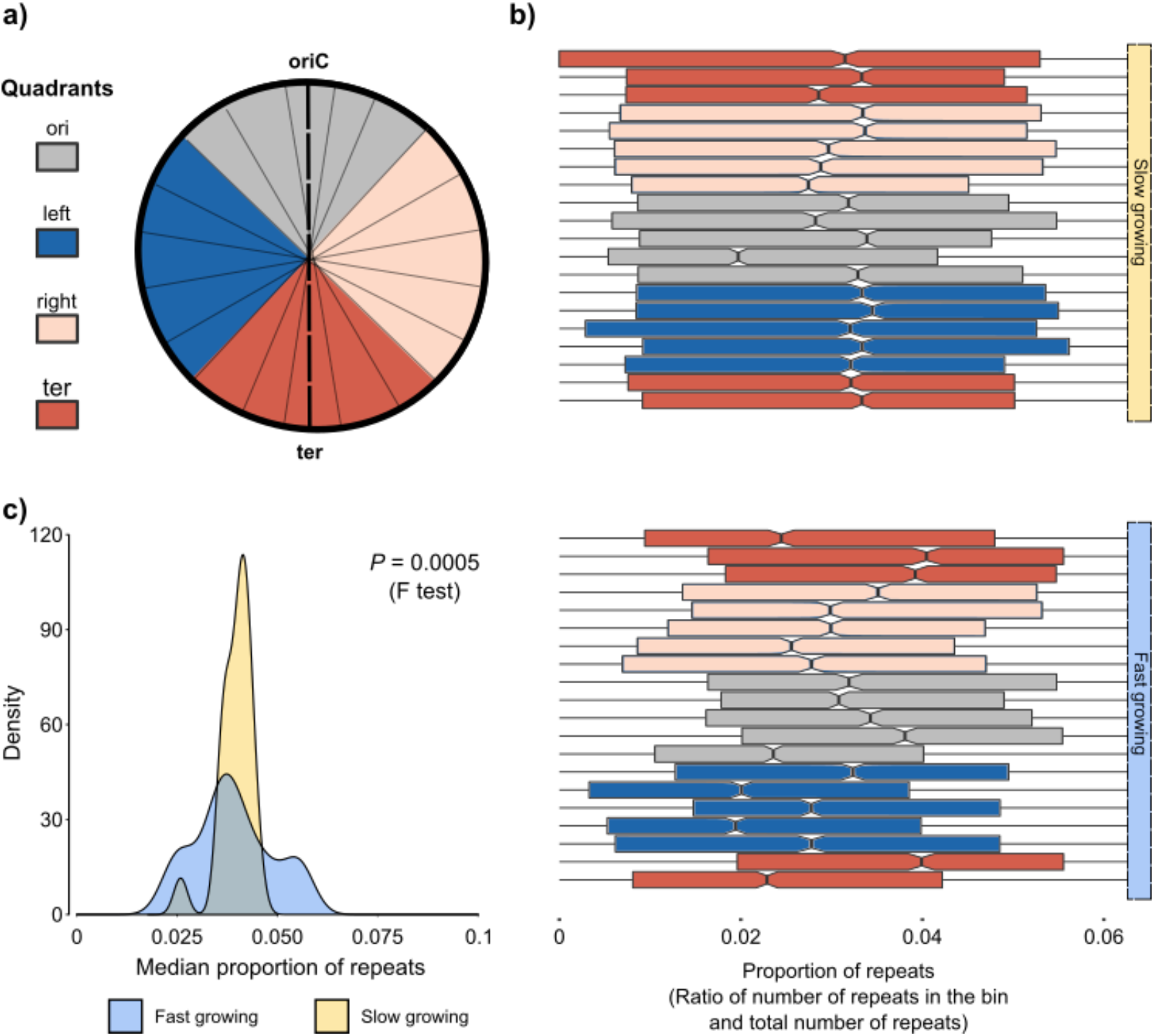
(a) Cartoon picture representing bacterial chromosome with oriC-ter axis and 20 bins. The colours represent different quadrants made using oriC and ter as reference. (b) Boxplots of repeat proportions in bins across bacterial genomes. The colours represent different quadrants made using oriC and ter as reference. The upper plot represents the distribution in slow growing bacteria while the lower set of the plot represent distribution in fast growing bacteria. (c) Kernel density plot of median proportion of repeats in each bin from (d) in fast and slow growing bacteria. F test using var.test() function in R is used to calculate the difference in variance in the two distributions and obtain the *P* value.

### oriC proximal repeats are involved in lower long-range interactions

To identify patterns inthe repeat proximity to oriC and ter, we divided the genome into four quadrants as described earlier in (Khedkar & Seshasayee 2016). The ori quadrant is centred around oriC and the ter quadrant is around ter. The quadrants between oriC and ter on either replichores are called right and left quadrants (Figure 5a). For each quadrant pair, we calculated the proportion of repeat pairs as the count of repeat pairs in the pair of quadrants to the total number of repeat pairs in that genome. We observed that repeat pairs across genomes are present closer to each other i.e., as intra-quadrant pairs (both repeats of a pair in the same quadrant) as compared to inter-quadrant pairs (repeats of a pair in different quadrants). This is clearly evident from the ‘X’-shaped pattern in the heatmap of median repeat pairs proportion across bacteria (Figure 5b).

**Figure 5.**
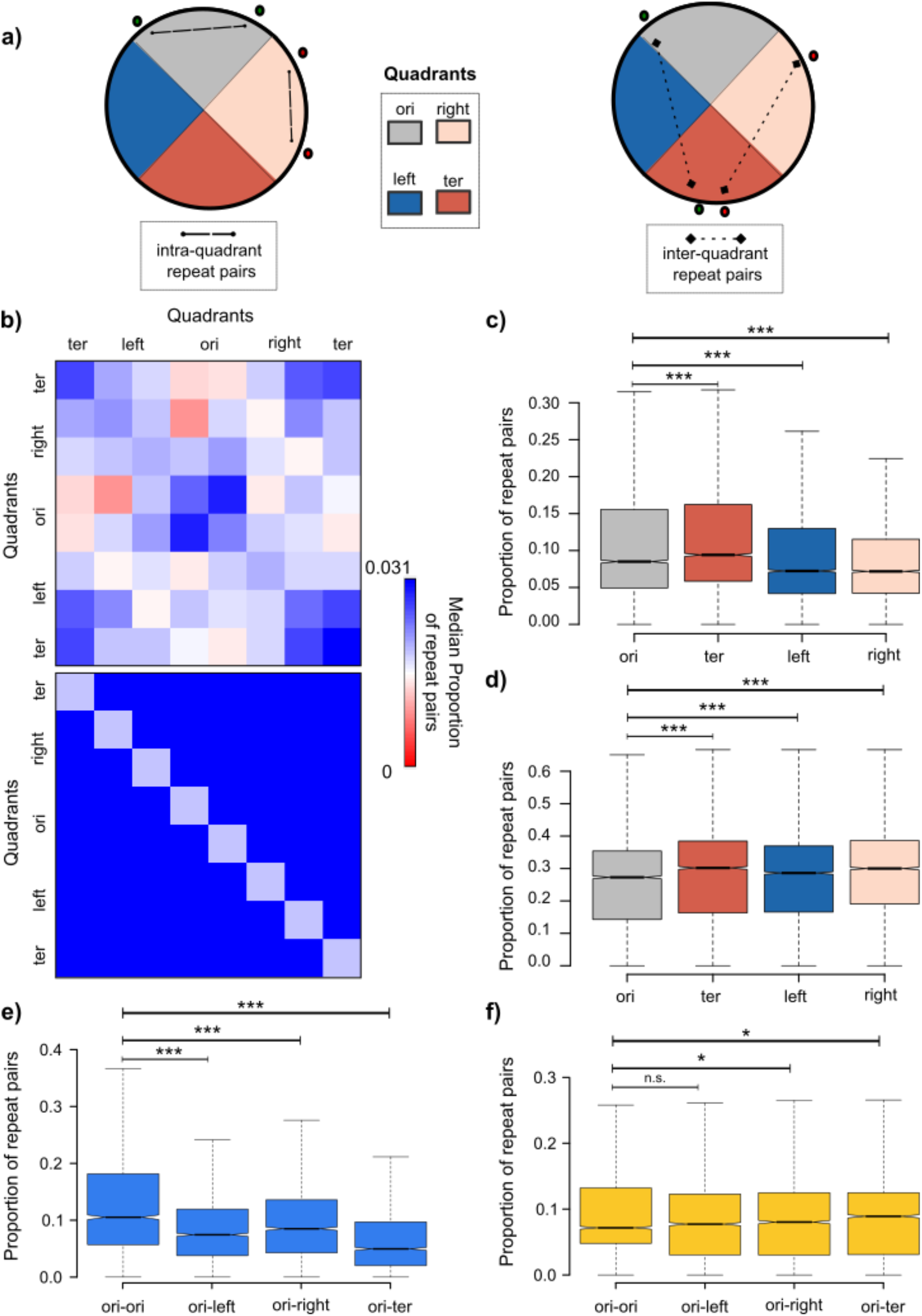
(a) Cartoon picture representing bacterial chromosome divided into four quadrants. The upper picture shows intra-quadrant repeat pairs (dashed line with dots at the end) i.e.; both repeats of a pair are in the same quadrant while the lower picture shows inter-quadrant repeat pairs (dotted line with diamonds at the end) i.e.; repeats of a pair are in different quadrants. The colours represent different quadrants made using oriC and ter as reference. (b) Heatmap of median (across genomes) proportion of repeat pairs across bin pairs. ‘X’-shaped pattern is visible indicating a higher proportion of intra-quadrant repeat pairs. Boxplots of proportion of repeat pairs present as (c) intraquadrants pairs and (d) inter-quadrants pairs across quadrants. Boxplots of proportion of repeat pairs present as inter-quadrant pairs with ori as one of the quadrants in (e) fast growing bacteria and (f) slow growing bacteria Wilcoxon’s rank-sum test is used to calculate the significance in the differences of two medians in each pair of comparison. (n.s. :P>= 10-2, * : P < 10-2, ** : P< 10-4, *** : P< 10-6)

On comparing repeat pairs proportion across quadrants, we observed that both ori and ter quadrants show the highest proportion of intra-quadrant repeat pairs (Figure 5c). Incontrast, we found that the proportion of all inter-quadrant repeat pairs involving ori quadrant (one repeat of a pair in ori quadrant and the other repeat in any of the left, right or ter quadrant) is least when compared to all other quadrants (Figure 5d). Furthermore, we observed that this difference in proportion of inter-quadrant repeat pairsand intra-quadrant pairs involving ori is more significant in fast growing bacteria as compared to the slow growing bacteria (Figure 5e and 5f).

Thus, despite having more repeats near the origin of replication, fast-growing bacteria lack repeats potentially involved in long-range recombination events involving oriC-proximal regions that can substantially disrupt the replicative structure.

### Distribution of repeats across replichores is different on the basis of their orientation

The impact of repeat pairs on fitness would depend on the repeat type and on whether the two members of the pair are present on the same replichore or on alternative replichores. Towards studying this effect, we categorised repeat pairs into two categories; inter-replichore repeats i.e., repeat pairs across two halves or replichores divided by the oriC-ter axis and intra-replichore repeats as repeat pairs present on the same side of the axis (Figure 6a). This is of particular interest to inverted repeats: for example, inversions caused by recombination between repeats in the same replichore switch affected genes from leading to lagging strand and vice-versa whereas that between repeats across replichores do not. Additionally, inversion of the gene also leads to change in the gene dosage due to the relative change in oriC proximity in both inter- and intra-replichore pairs; however, if the two recombining inter-replichore inverted repeat elements are positioned symmetrically around the oriC, the inversions would affect gene dosage minimally. In case of deletions or duplications mediated by direct repeat pairs, the gene dosage of the affected region will be altered similarly in both intra-replichore and inter-replichore arrangements. However, deletion of the oriC itself would be immediately lethal and large duplication across replichores mediated by direct repeats would be unstable.

**Figure 6.**
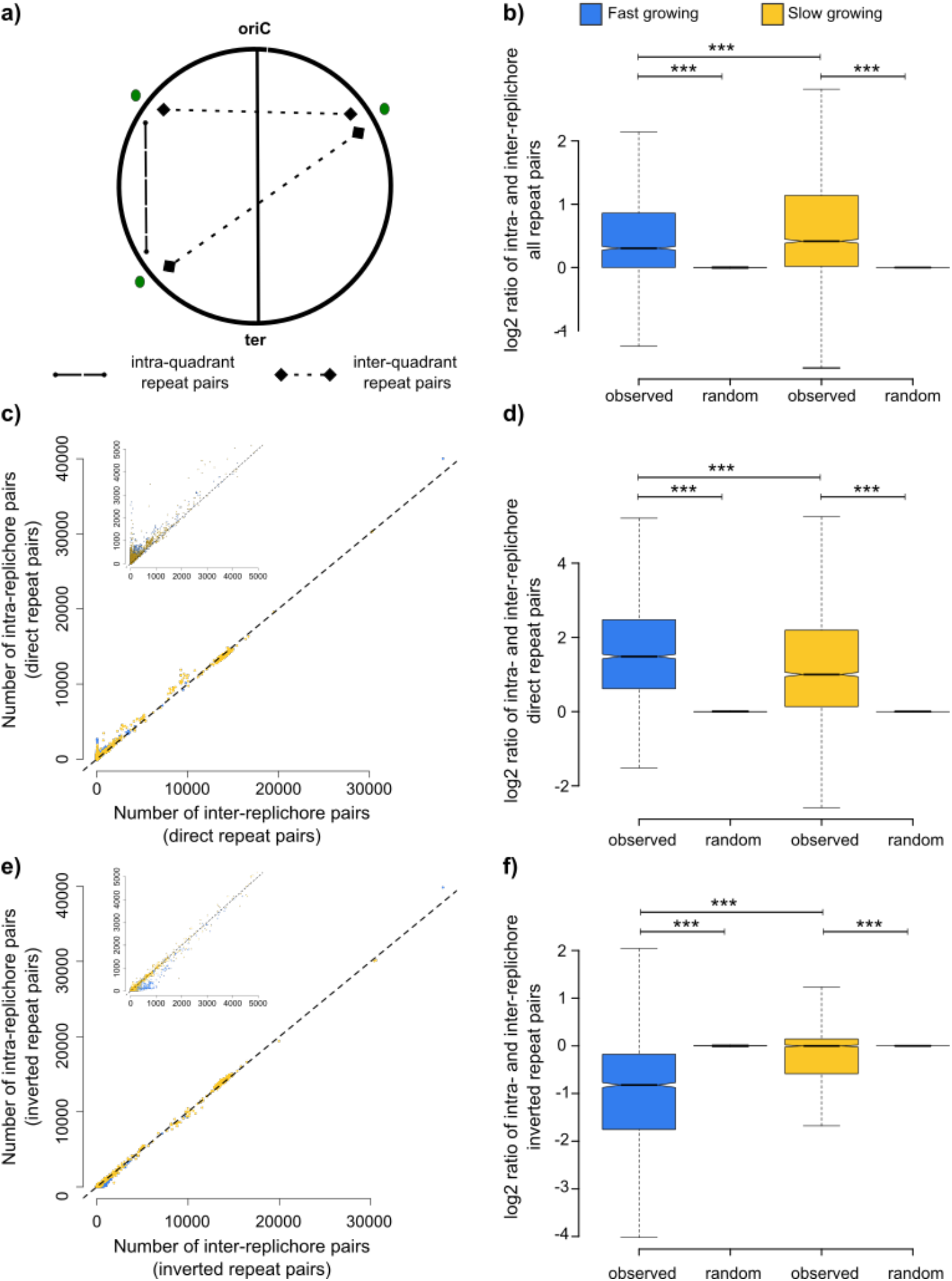
(a) Cartoon picture representing bacterial chromosome divided into two replichores across oriC-ter axis. Green dots represent repeats present as intra-replichore pair (dashed line with dots at the end) and as inter-replichore pair (dotted line with diamonds at the end). (b) Boxplots of proportion of intra-replichore repeat pairs shown as log2 (number of intra-replichore repeat pairs / number of inter-replichore repeat pairs) in all repeat pairs (both direct and inverted repeats). (c) Scatter plot of number of inter- and intra-replichore direct repeat pairs with coloured dots indicating bacterial growth categories (blue: fast growing, yellow: slow growing). (d) Boxplots of proportion of intra-replichore repeat pairs shown as log2 (number of intra-replichore repeat pairs / number of inter-replichore repeat pairs) for only direct repeat pairs in fast growing and slow growing bacteria. (e) Scatter plot of number of inter- and intra-replichore inverted repeat pairs with coloured dots indicating bacterial growth categories (blue: fast growing, yellow: slow growing). (f) Boxplots of proportion of intra-replichore repeat pairs shown as log2 (number of intra-replichore repeat pairs / number of inter-replichore repeat pairs) for only inverted repeat pairs in fast growing and slow growing bacteria. Wilcoxon’s rank-sum test is used to calculate the significance in the differences of two medians in each comparison between fast and slow growing bacteria whereas Wilcoxon’s signed-rank test (paired) is used to calculate the significance in the differences of medians of the observed and random populations in each comparison. (n.s. :P>= 10-2, * : P < 10-2, ** : P< 10-4, *** : P< 10-6)

To perform statistical inference on the proportion of intra/inter-replichore repeats, we randomly allotted a replichore to all the repeat elements present in the genome. We then calculated the proportion of intra-replichore repeats in that randomisation event. For every genome, a median of the proportions obtained from 1000 randomisation events was taken. On comparing the distribution of observed genomic proportions of intra-replichore repeats to the median proportions obtained by randomization, we found that repeat pairs are present significantly higher (Wilcoxon signed rank test (paired); P< 10^−10^) in the same replichore (intra-replichore) as compared to that by random event, in both fast and slow growing bacteria (Figure 6b). This proportion is significantly less in fast growing bacteria as compared to slow growing bacteria. (Wilcoxon rank sum test; P < 10^−10^) (Figure 6b).

On comparing different orientations of repeat pairs with reference to the replichores, we found that direct repeats were found higher than by random chance in intra-replichore arrangement (Figure 6c and 6d) (Wilcoxon signed rank test (paired); P< 10^−10^) (Figure 6e and 6f), whereas inverted repeat pairs were present more in inter-replichore arrangement (Wilcoxon signed rank test (paired); P< 10^−10^) (Figure 6e and 6f). When compared between the two bacterial growth categories, we observed that the number of intra-replichore direct repeats was significantly higher in fast growing as compared to slow growing bacteria (Wilcoxon rank sum test; p-value = 10^−10^) whereas intra-replichore inverted repeat pairs were found significantly less frequently in fast-growing organisms (Wilcoxon rank sum test; P< 10^−10^) (Figure 6d and 6f).

### Genomic distribution of repeats enables minimal disruption to the replication dependent genome organisation

If repeat pairs are present intra-replichore, a higher spacing between the pairs would lead to the rearrangement of a larger genomic region and thus disrupt the gene dosage gradient to a greater extent. On the other hand, if inter-replichore repeats are present symmetrical about the oriC-ter axis, the extent of disruption to gene dosage would be minimal.

In case of direct repeat pairs, we observed that the average distance between repeat pairs in the same replichore is significantly less (Wilcoxon signed rank test (paired); P< 10^−10^) than the average distance obtained from the randomly shuffled positions of the repeats in both fast and slow growing bacteria. This can be seen from the darker diagonal in the heatmap of median proportion of direct repeats (Figure 7a). This distance was significantly lower (Wilcoxon rank sum test; P = 7.7*10^−6^) in fast growing bacteria when compared to that in slow growing bacteria (Figure 7b). Surprisingly, in case of inverted repeat pairs, the diagonal in the heat map of median proportion of inverted repeats is not visible (Figure 7c). The distance between repeats is significantly higher than random (Wilcoxon signed rank test (paired); P < 10^−10^) with no difference in fast and slow growing bacteria. (Figure 7d).

**Figure 7.**
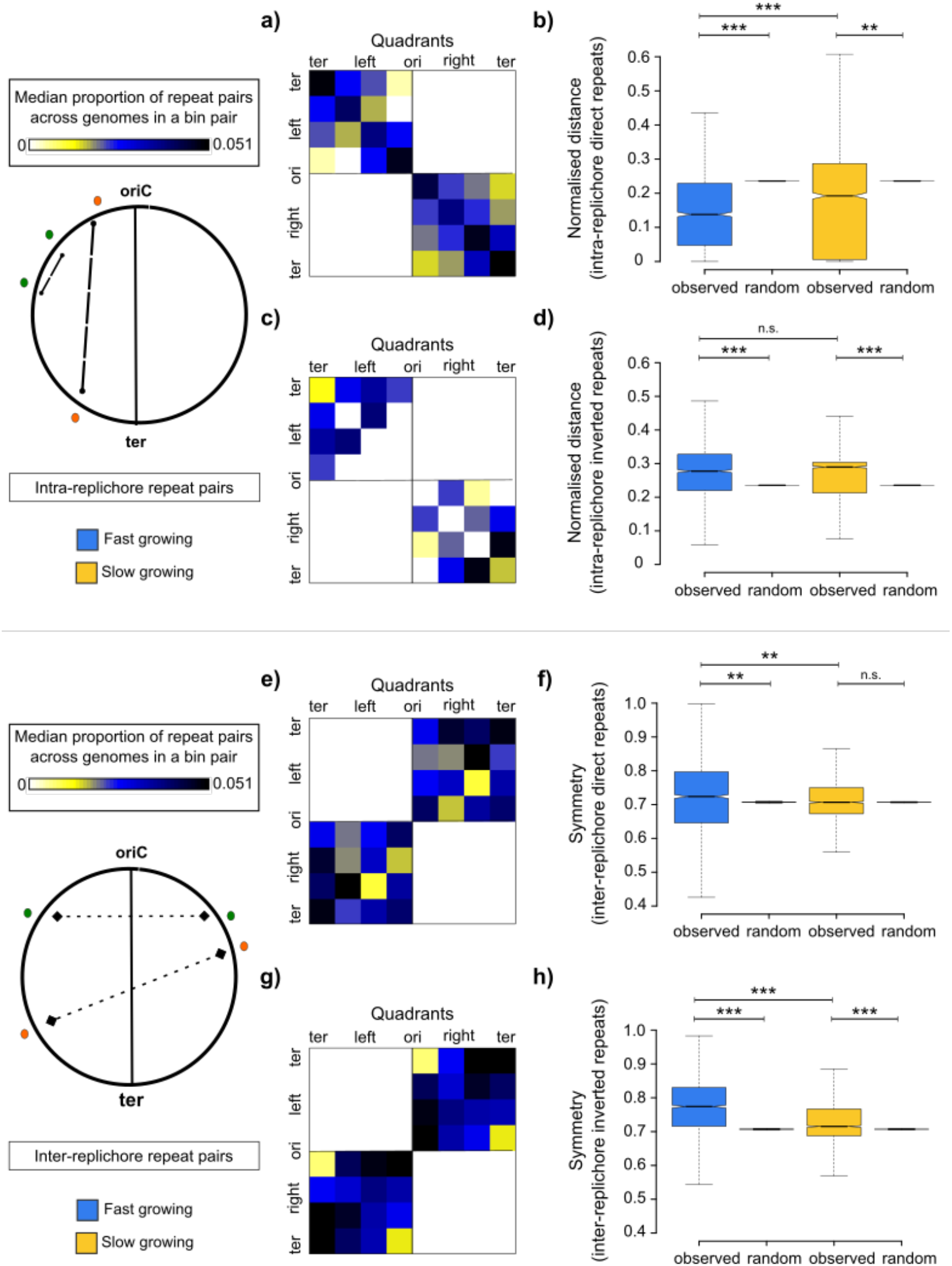
(a) Heatmap of median (across genomes) proportion of intra-replichore direct repeat pairs across bin pairs. (b) Boxplots of normalised distances between intra-replichore direct repeat pairs in fast and slow growing bacteria. (c) Heatmap of median (across genomes) proportion of intra-replichore inverted repeat pairs across bin pairs. (d) Boxplots of normalised distances between intra-replichore inverted repeat pairs in fast and slow growing bacteria. (e) Heatmap of median (across genomes) proportion of inter-replichore direct repeat pairs across bin pairs. (f) Boxplots of symmetry between inter-replichore direct repeat pairs in fast and slow growing bacteria. (g) Heatmap of median (across genomes) proportion of inter-replichore inverted repeat pairs across bin pairs. (h) Boxplots of symmetry between inter-replichore inverted repeat pairs in fast and slow growing bacteria. Wilcoxon’s rank-sum test is used to calculate the significance in the differences of two medians in each comparison between fast and slow growing bacteria whereas Wilcoxon’s signed-rank test (paired) is used to calculate the significance in the differences of medians of the observed and random populations in each comparison. (n.s. :P>= 10-2, * : P < 10-2, ** : P< 10-4, *** : P< 10-6)

To find out if repeat pairs are present symmetrical around oriC, we calculated the symmetry of inter-replichore repeat pairs as 1 - absolute difference in the normalised relative position of the repeat units with respect to oriC. These values theoretically range from 0 to 1 with absolute difference of ‘1’ implying that the repeat pair are symmetrical around oriC while a repeat pair which is most asymmetric around the oriC-ter axis i.e., present on the axis itself, has a value of 0. While direct repeats are slightly more symmetric than random in terms of statistical significance, the effect size is small (Figure 7e-7h). This significance holds only for fast-growing, but not in slow growing bacteria (Wilcoxon signed rank test (paired); P< 3.17*10^−5^ for fast growing bacteria; Figure 7f). However inverted repeat pairs are present symmetrical in both fast and slow growing bacteria with a clear difference from random chance (Wilcoxon signed rank test (paired); P< 10^−10^) and these repeats were significantly more symmetrical in fast growing bacteria (Wilcoxon rank sum test; P< 10^−10^) (Figure 7h).

Our observations of symmetric presence of long DNA repeats are consistent with previous reports on symmetric inversions (‘X’ shaped patterns on pairwise genome alignment) in *Azotobacter vinelandii* and *Vibrio chloreae* genomes (Repar & Warnecke 2017) and symmetric translocations in *Caulobacter crescentus* (Khedkar & Seshasayee 2016). In ascatter plot of normalised positions of inter-replichore repeat pairs, we found that most of the repeats in *Vibrio* and *Caulobacter* are closer to the diagonal which indicate their symmetry around oriC as seen in (Figures 8a and 8b). This can be clearly seen in the distribution of absolute difference in the relative positionsof the repeats, where the mode is closer to 0 (closer / symmetric the repeat pairs, lower the difference in their relative positions). Though we don’t see a such clear symmetry in the position of repeats with respect to the diagonal in *Azotobacter*, the distribution of absolute difference in positions has mode closer to 0 (Figure 8c). These examples also show closeness between intra-replichore repeat pairs.

**Figure 8.**
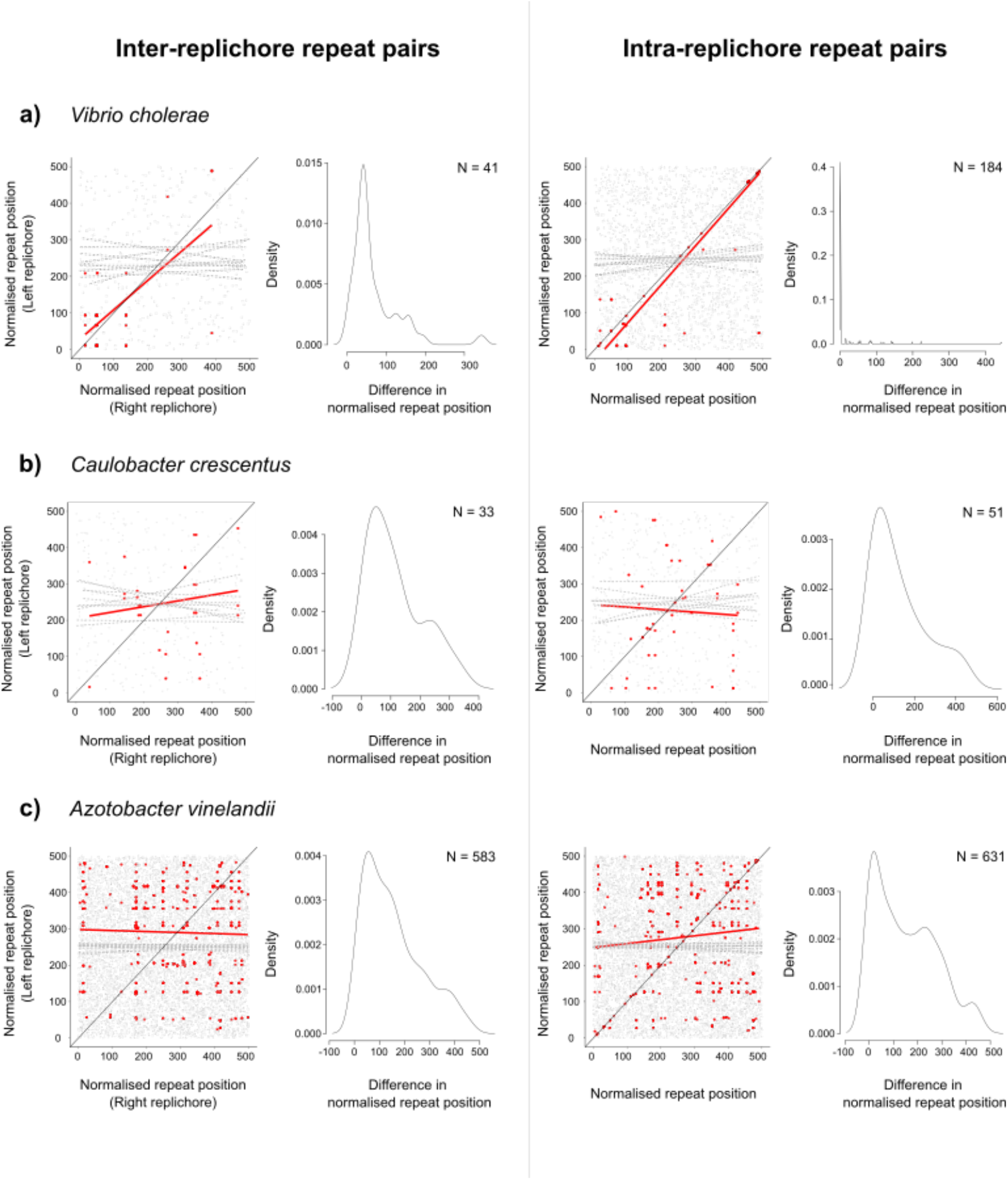
Scatter plot of normalised repeat positions of the repeats in a pair in case of inter-replichore and intra-replichore repeat pairs with kernel density distribution of absolute differences in the normalised relative positions of the repeats in (a) *Vibrio cholerae*, (b) *Caulobacter crescentus*, and (c) *Azotobacter vinelandii*. The red coloured points in the scatter plot represents the positions of observed dataset with red line indicating the linear model fit whereas grey points and dotted lines represent the positions of randomly shuffled set for 10 iterations.

Long DNA repeats being closer to each other and symmetrical around oriC imply that the genomic organisation of repeat pairs facilitates minimal disruption to the replication associated genome organisation with more closeness and symmetry in fast growing bacteria.

## Discussion

In this study, using fully available bacterial genomic sequences, we have identified long DNA repeats, which are potential players in intra-chromosomal rearrangements in bacteria. We found that these repeats are present in a non-random manner across bacteria of different genera and classes. In our exploration of replication dependent constraints on genome organisation of these repeats, we found that these repeats are non-randomly distributed in terms of their overall density, orientation, proximity to oriC, and with respect to spacing between them as well as their symmetry around the oriC-ter axis. Taken together, in fast growing bacteria, where the effect of replication on genome organisation is strong, long DNA repeats are organised in a way that results in a relatively minimal disruption to the replication dependent genomic architecture. Our study reinforces the idea that replication is a prominent factor imposing selection on genomic organisation and therefore the positioning of long DNA repeats in these genomes.

### Repeat-mediated structural variations

Long intra-chromosomal DNA repeats are statistically unlikely to be present in a bacterial genome (Rocha et al. 1999). Despite this, bacteria across genera and classes are abundant in such repeats. On homology-based recombination, long repeats lead to deletion or duplication of the genomic segment or inversion of the region flanked by them. Laboratory evolution studies from various groups in different genomes of *Escherichia, Bacillus*, and *Salmonella* have shown that these structural variations affect bacterial fitness in different stresses and can be adaptive in nature under different conditions (Veetil et al. 2020; Srinivasan et al. 2015; Rocha et al. 1999; Sonti & Roth 1989; Maharjan et al. 2013; Achaz et al. 2003). Different types of structural variations in the genome impact bacteria differently. Deletions or duplications predominantly affect gene dosage while inversions change gene orientation. Our observation of lower proportions of inverted repeats suggests a selection against inversions in the genomes. This underrepresentation of inverted repeats is prominent in genomes that have low number of repeats and its proportion eventually plateaus with increase in the total number of repeats. This observation is consistent withand strengthens the findings by (Achaz et al. 2003).

### Rearrangements and genome organisation

Bacterial genomes exhibit organisational features like strand skewness of essential genes and oriC proximity of highly expressed genes (Rocha 2004; Couturier & Rocha 2006; Rocha 2003). Any change to the organisation carries some cost and thus affects fitness. Changes in the position of a gene can lead to changes in their dosage and thus expression; this might further hamper bacterial growth (Bryant et al. 2014). Alternatively, any shift in the position of the oriC or a disruption of oriC functioning, can impact gene expression on a global scale and might be substantially deleterious for the bacteria (Veetil et al. 2020; Dimude et al. 2018; Bryant et al. 2014; Duigou & Boccard 2017). Changes in gene orientation, caused by inversions, might result in head-on collisions of DNA and RNA polymerase. This effect, which is more likely to happen at highly expressed genes, might further cause DNA damage and inhibit growth consequently.

Changes in genome organisation in terms of orientation and positions relative to oriC affect bacterial fitness and thus are under selection. Likewise, anything mediating these changes will also be under selection. Since these changes are often caused by repeat-mediated rearrangements, this begs the question on how these repeats are distributed in the genome. We show that long repeats are distributed in a non-random manner across bacteria. This observation is in support of Repar & Warnecke 2017 who showed non-random inversion landscapes in bacterial and archaeal genomes. Though regions near oriC and ter are repeat-rich in fast growing bacteria, these are positioned in a manner that they cause short-range rearrangements and thereby disrupt genome organisation minimally. This was supported by the study by (Valens et al. 2016) demonstrating lower inversion frequency between ori and right macrodomains relative to intra-macrodomain frequency.

Previous reports have found symmetry in the inversions observed across bacterial genomes (Helm et al. 2003; Repar & Warnecke 2017; Eisen et al.; Kong et al. 2009). Consistent with this, we find that inter-replichore inverted repeat pairs tend to be more symmetric around the oriC-ter axis than expected by random chance. Though inverted repeats are less prevalent than direct repeats, they tend to occur inter-replichore. Recombination between inverted repeat pairs intra-replichore can affect genome organisation by changing gene orientation and thus dosage. Across replichores, symmetrically positioned inverted repeat pairs will minimise disruption to genome organisation. Direct repeat pairs, often intra-replichore, are present closer to each other compared to inverted repeat pairs. Though symmetry of inter-replichore direct repeat pairs is significantly more than random, the magnitude of the difference in median values is small. The statistical significance in this case is likely due to large sample size, and disappears when we consider a single random representative genome per species for analysis (Figure S2). These observations of close intra-replichore repeats and symmetric inter-replichore repeats are more prominent in fast growing bacteria when compared with slow growing bacteria and hold true even after removing species redundancy (Figure S2) and in most of the comparisons in major phyla (Figure S3) with varying degrees of significance.

Since selection would act on rearrangements and these are a function of not only repeat pair presence and distance / symmetry but also their propensity to interact physically, variations in 3D conformations in the chromosome can enhance, limit or even abolish selection on repeat positions. We took 3C or Hi-C data available for *C.crescentus, E.coli, B.subtilis*, and *M. pneumoniae* from previous studies (Le et al. 2013; Lioy et al. 2018; Wang et al. 2014; Trussart et al. 2017) and compared the normalised interaction scores (as reported in the study) of the genomic regions (bin pairs) containing repeats to the regions devoid of repeats in the WT condition. Though we see a significant difference in interaction scores between the groups in *E. coli*, we do not see much difference in the average interaction scores of *C. crescentus* and *B. subtilis*. The average interaction scores of regions with repeats were significantly lower with rest of the regions in *M. pneumoniae* (Figure S4). The inconsistency in patterns observed across these few limited bacterial genomes suggests that the product of repeat position and repeat interaction is likely to vary considerably across organisms.

### Repeat distribution patterns: mechanistic processes and selection

How do repeats originate? (Achaz 2002) propose that repeats arise at the first instance by tandem duplication and such repeat pairs are direct repeats. Additional direct repeats and inverted repeats are created by other events causing structural rearrangements such as inversions, translocations, inversions, etc. We performed a toy simulation in which a single repeat element placed in one of 100 random genomic slots is allowed to undergo several tandem duplications and rearrangements at arbitrary rates; in this simulation, we assume that rearrangements maintain or reverse the orientation of the affected repeat at equal probability. By definition this would produce more direct than inverted repeats as observed in real data, unless the rate of rearrangements is high enough and the probability of rearrangements generating inverted repeats is much higher than those producing more direct repeats.

As expected, even without invoking selection, this model of repeat generation would place direct repeats closer to each other than expected by pure random chance. If selection were imposed over this such that the probability of the loss of an intra-replichore direct repeat is higher the farther the members of the pair are from each other, the distribution of the distance between the repeat elements would decrease (Figure S5a for direct repeats). Our observation from genome data in which intra-replichore direct repeats are closer together more in fast-growing than in slow-growing bacteria suggests a role for selection, unless this difference could be explained purely by more frequent replication events causing more tandem duplications. Inter-replichore direct repeat pairs would almost always be generated by translocation-related processes and no additional origin of selection, beyond distance between them, can be envisaged for them.

Inverted repeats present a more interesting situation. They are not generated by tandem duplication and assuming that translocation events place them at random positions on the genome, there is no reason to assume that without selection their relative position should be any different from random. Curiously, we observe in bacterial genomes that intra-replichore inverted repeats are placed further apart from each other than random expectation (though this trend is lost when we picked one random representative genome per species). Assuming this pattern is biologically meaningful, at least in a subset of genomes considered here, there are two possibilities that can cause such a pattern to arise: (a) there could be a minimum distance over which translocations can occur in some genomes, and this would likely push inverted repeat pairs further apart than expected by random chance; (b) selection operates *against* inverted repeat pairs that are relatively close to each other (Figure S5a for inverted repeats), presumably because chromosome conformations make recombination events between closely positioned repeats more likely; this would be less effective for direct repeat pairs for which the process of generation by tandem duplication would be a strong counterforce. Inter-replichore inverted repeat pairs would be positioned randomly in the absence of selection. It would be reasonable to assume that selection would operate against inter-replichore inverted repeat pairs that are positioned asymmetrically about the oriC-ter axis. Therefore, imposing a probability of repeat loss under selection that is inversely proportional to symmetry would tend to place inter-replichore inverted repeat pairs in more symmetric positions than random (Figure S5b) producing a distribution of repeat pair symmetrythat is shifted to the right, similar to the results obtained here as well as for the translocation data observed by (Khedkar & Seshasayee 2016).

Further exploration of this toy model with deep sampling of the parameter space might provide additional insight into the processes underlying the repeat distributions observed in this study.

## Conclusion

Maintenance of genome organisation and chromosomal variations are inter-dependent and go hand in hand in the course of genome evolution. On one hand, structural variations play a major role in adaptation and evolution of the organism (Straus & Hoffmann 1975; Sonti& Roth 1989; Rocha et al. 1999; Rocha 2002; Srinivasan et al. 2015; Veetil et al. 2020; Gorkovskiy & Verstrepen 2021), and on the other hand, theycan disrupt genomic organisation and thus affect fitness (Hill & Harnish 1981; Hill & Gray 1988; Rocha 2004; Adler et al. 2014). Taken together, byanalysing ~6k genomes, our study finds association between intra-chromosomal long DNA repeats and replication-dependent bacterial genome organisation. However, we considered only homologous recombination as the mechanism of repeat-mediated disruptions and do not take account of disruptions mediated by transposition events. We suggest maintenance of replication dependent genome organisation as a selection pressure in the positioning of long DNA repeats, involved in causing chromosomal rearrangements. There exists a balance between the interplay of genome variability caused by repeats and genome maintenance, and the presence of repeats at certain positions enables reduction in disruption to the replication-dependent genome organisation.

## Acknowledgment

We thank Anjana Badrinarayanan, Dasaradhi Palakodeti and Dimple Notani for helpful discussions as part of the thesis advisory committee. We also thank Akshara Dubey, Meghna Nandy, Mohak Sharda and other members of the ASN lab for valuable inputs and discussions throughout the study. We appreciate contributions by Akshaya Seshadri, Anurag Kumar Singh, Kushi Anand and Neha Sontakke for proofreading the manuscript. This work is supported by a DBT/Wellcome Trust India Alliance Intermediate Fellowship (IA/I/ 16/2/502711 to A.S.N.S.), and core funding from the Department of Atomic Energy, Government of India (Project No. 12-R&D-TFR-5.04-0800).

## Data availability

All relevant data and scripts for analyses are available on Figshare via https://tinyurl.com/2b2e5h6s.

## Supplementary Figures

**Figure S1.**
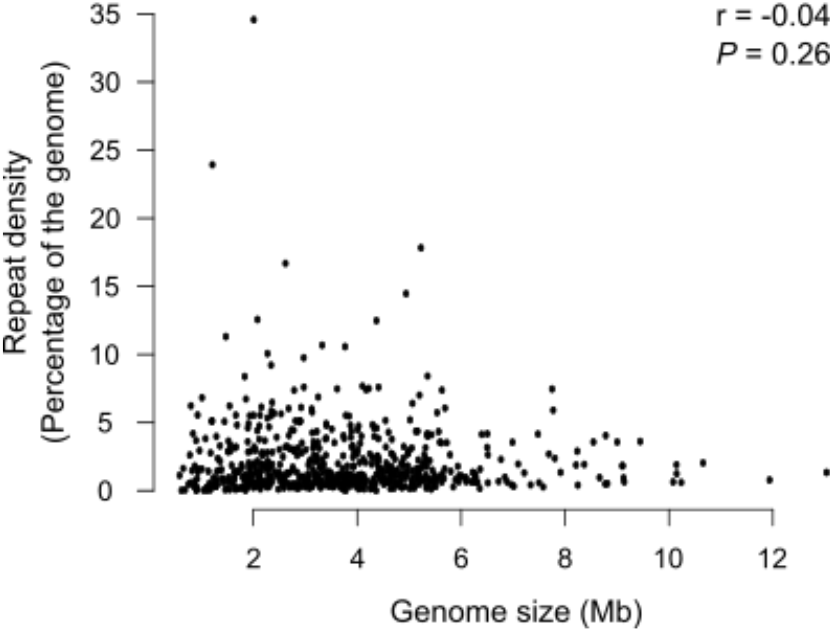
Scatter plot showing relation between repeat density and genome size (Megabases; Mb). The correlation is calculated by Pearson’s correlation using cor.test() function in R. The genomes here are selected randomly to remove species redundancy i.e., one species is represented bya single genome. The correlation is calculated by Pearson’s correlation using cor.test() function in R. N (Total number of genomes) = 653

**Figure S2.**
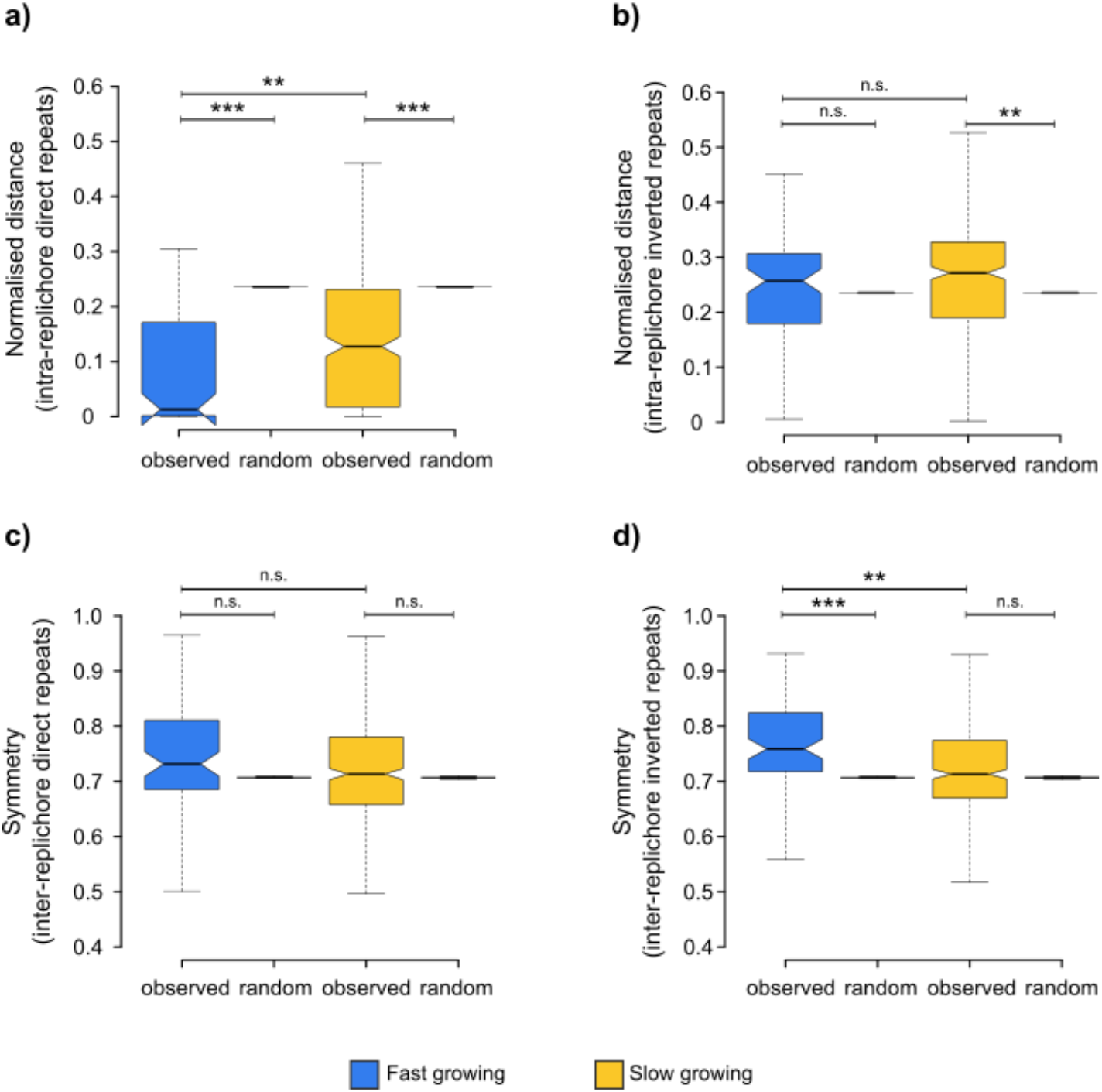
Boxplots of normalised distances between intra-replichore(a) direct repeat pairs and (b)inverted repeat pairs in fast and slow growing bacteria. Boxplots of symmetry between inter-replichore replichore (c) direct repeat pairs and (d) inverted repeat pairs in fast and slow growing bacteria (as in Figure 7). The genomes here are selected randomly to remove species redundancy i.e., one species is represented by a single genome. Wilcoxon’s rank-sum test is used to calculate the significance in the differences of two medians in each comparison between fast and slow growing bacteria whereas Wilcoxon’s signed-rank test (paired) is used to calculate the significance in the differences of medians of the observed and random populations in each comparison. (n.s. :P>= 10-2, * : P < 10-2, ** : P< 10-4, *** : P< 10-6)

**Figure S3.**
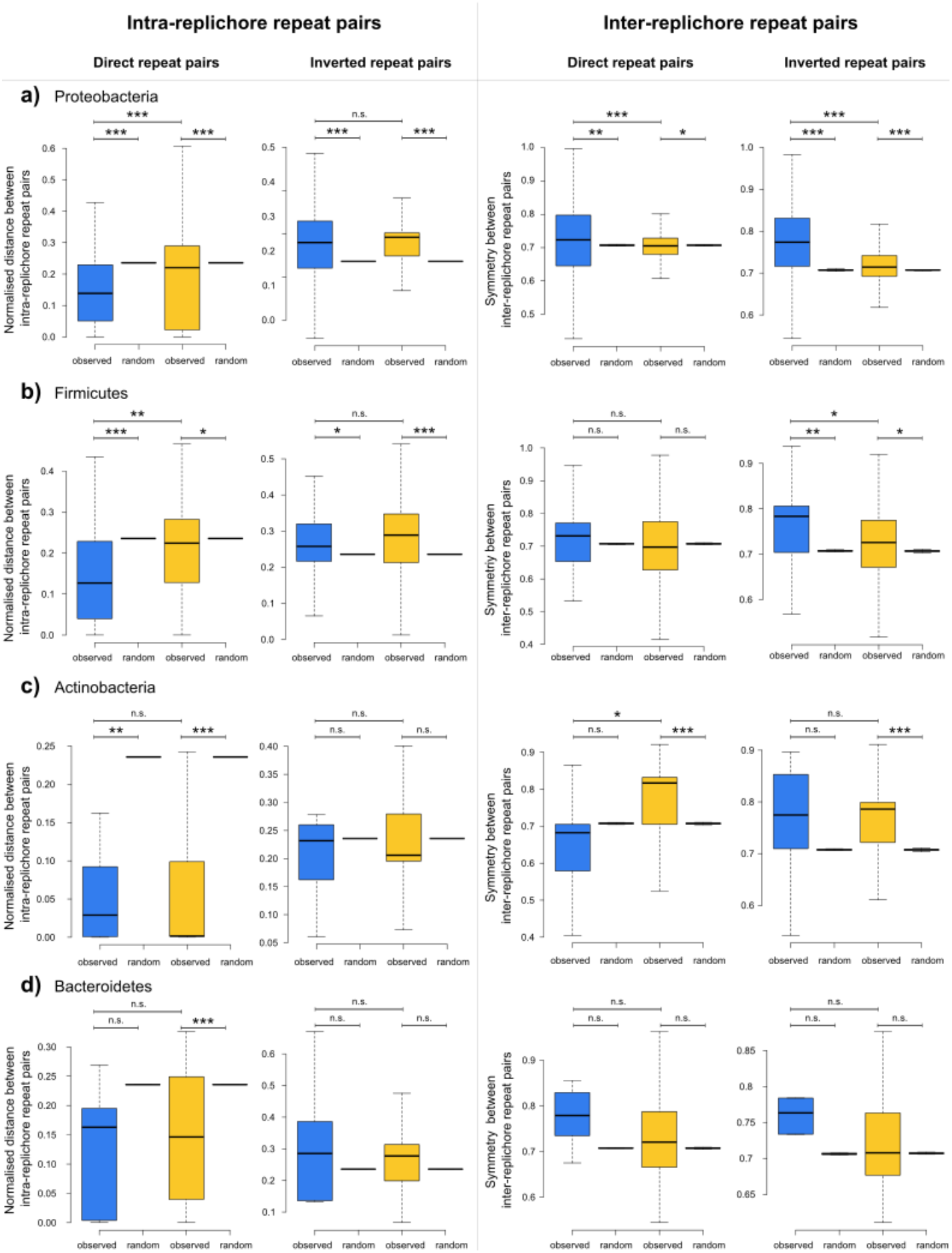
Boxplots of normalised distances between intra-replichore direct and inverted repeat pairs and symmetry between inter-replichore pairs in fast and slow growing bacteria (as in Figure 7) in (a) Proteobacteria, (b) Firmicutes, (c) Actinobacteria, and (d) Bacteroidetes. Wilcoxon’s rank-sum test is used to calculate the significance in the differences of two medians in each comparison between fast and slow growing bacteria whereas Wilcoxon’s signed-rank test (paired) is used to calculate the significance in the differences of medians of the observed and random populations in each comparison. (n.s. :*P*>= 10^−2^, * : *P* < 10^−2^, ** : *P*< 10^−4^, *** : *P*< 10^−6^)

**Figure S4.**
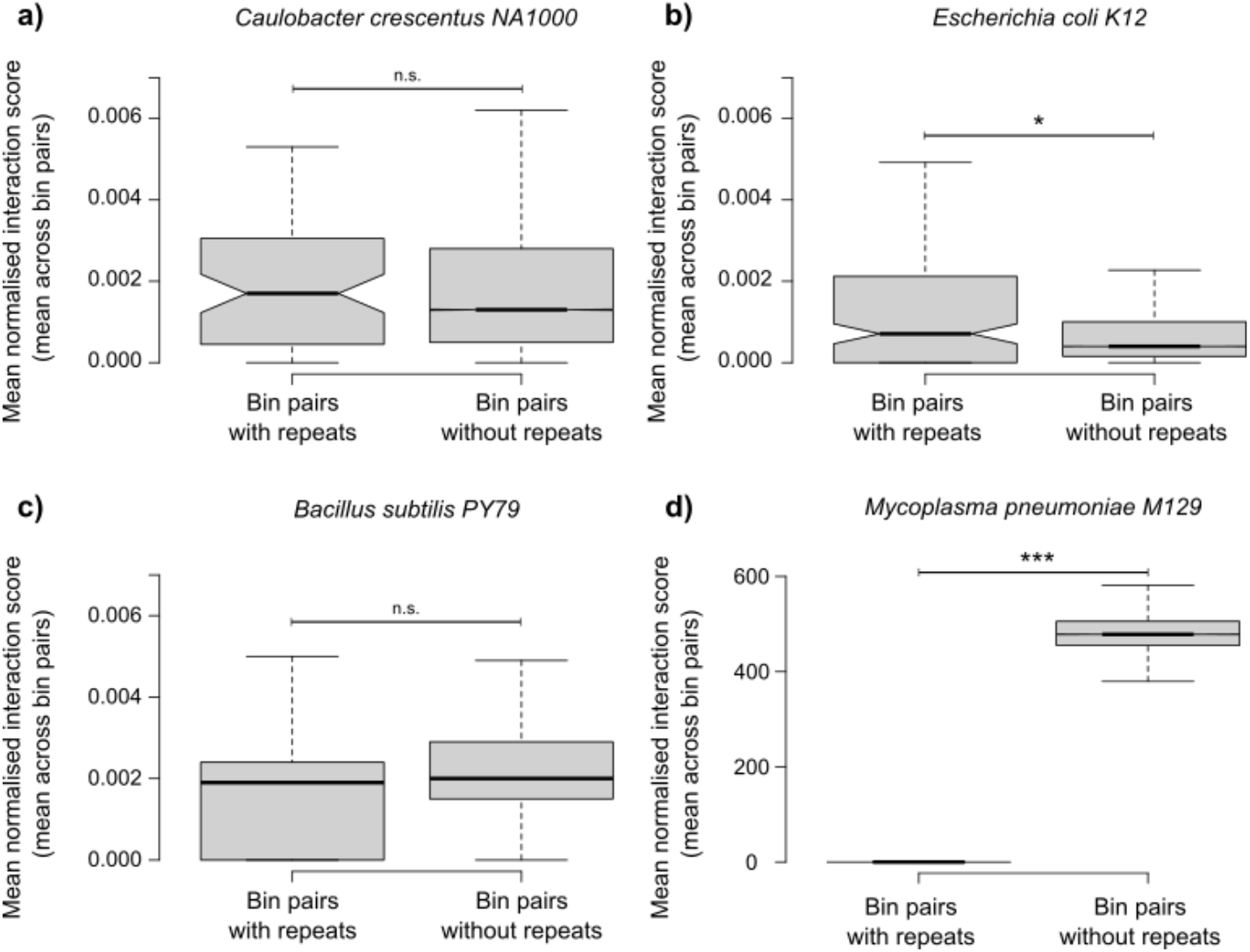
Boxplots of mean normalised interaction score (e.g., mean normalised interaction score for repeats in bin A and B = (normalised interaction score for pair AB = normalised interaction score for BA)/2) for bins containing repeat pairs and bins without repeat pairs in (a) *Caulobacter crescentus NA1000* (bin size = 10 Kb), (b) *Escherichia coli K12*(bin size = 5 Kb), (c) *Bacillus subtilis PY79* (bin size = 10 Kb), and (d) *Mycoplasma pneumoniae M129* (bin size = 3Kb). (n.s. :*P*>= 10^−2^, * : *P* < 10^−2^, ** : *P*< 10^−4^, *** : *P*< 10^−6^)

**Figure S5.**
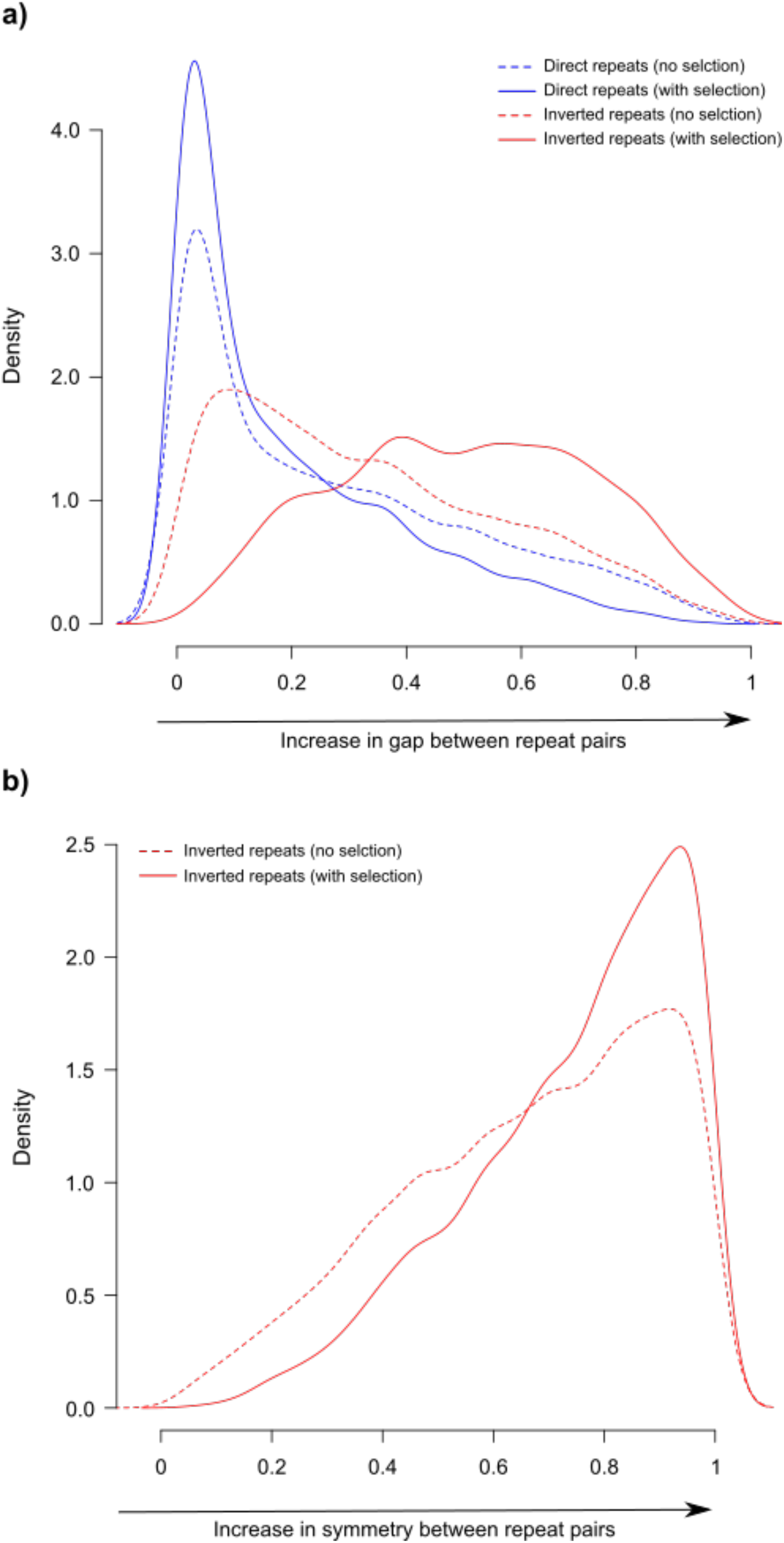
Kernel density plot showing (a) distribution of distance in intra-replichore repeat pairs and (b) symmetry in inter-replichore inverted repeat pairs. The dotted lines represent the distribution of distance / symmetry with no selection acting on the position of the repeats. The solid-coloured lines represent distributions obtained with selection acting on the genomic position of the repeats. The distributions are obtained after simulating generation of multiple copies of repeats over 10000 iterations, starting from a single copy, based on our interpretation of the model by (Achaz 2002)and applying distance / symmetry-based selection on these. For every repeat randomly positioned in one of 100 random slots, a probability of tandem duplication [chosen from a beta distribution β(0.05,2)] and translocation [chosen from β(0.2,2)] per iteration was allotted. Each tandem duplication event produces another copy of the duplicated repeat making the pair a direct repeat pair and places it in a position that is Poisson (1) slot away from the duplicated repeat. In case of translocation, the position of the new repeat was randomly picked from a uniform distribution (1-100) with equal probabilities of retaining or reversing the starting orientation. The process of duplication and translocation resulted in inter-and intra-replichore repeat pairs of both direct and inverted orientations. On average, each starting repeat produced ~23 pairs in this simulation. For direct repeat pairs, selection was imposed on repeat pairs in such a way that the probability of the loss of repeats would be higher with increase in gap between them. For inverted repeat pairs, the selection was imposed differently for intra- and inter-replichore repeats. For intra-replichore inverted repeats, the selection operates against inverted repeat pairs that are relatively close to each other (see Discussion on why this non-intuitive possibility was considered), whereas in case of inter-replichore inverted repeat pairs it would operate against repeat pairs that are positioned asymmetrically about the oriC-ter axis. Selection was implemented by a probability of a repeat pair being retained, the distance / symmetry described above being the mean of a beta distribution [β(5/(1-μ),5) or β (5,5(1- μ)/μ) depending on the direction of the tail].

